# A family of Tn7-like transposons evolved to target CRISPR repeats

**DOI:** 10.1101/2024.10.13.618069

**Authors:** Laura Chacon Machado, Joseph E. Peters

## Abstract

Tn7 family transposons are mobile genetic elements known for precise target site selection, with some co-opting CRISPR-Cas systems for RNA-guided transposition. We identified a novel group of Tn7-like transposons in Cyanobacteria that preferentially target CRISPR arrays, suggesting a new functional interaction between these elements and CRISPR-Cas systems. Using bioinformatics tools, we characterized their phylogeny, target specificity, and sub-specialization. The array-targeting elements are phylogenetically close to tRNA-targeting elements. The distinct target preference coincides with loss of a C-terminal region in the TnsD protein which is responsible for recognizing target sites when compared to closely related elements. Notably, elements are found integrated into a fixed position within CRISPR spacer regions, a behavior that might minimize negative impacts on the host defense system. These transposons were identified in both plasmid and genomic CRISPR arrays, indicating that their preferred target provides a means for both safe insertion in the host chromosome and a mechanism for dissemination. Attempts to reconstitute these elements in *E. coli* were unsuccessful, indicating possible dependence on native host factors. Our findings expand the diversity of interactions between Tn7-like transposons and CRISPR systems.

## Background

Transposons are mobile genetic elements that can move between locations within a genome. Tn7 and Tn7-like transposons are distinguished by their capacity to select specific targets for transposition (1). With these elements, transposition activity is latent until a preferred target is recognized. Tn7-like elements typically have two targeting systems, one that recognizes a conserved site in host chromosomes (homing) and a second that can recognize other mobile elements that allow transfer to new hosts, like conjugal/mobile plasmids (dissemination). These complementary pathways facilitate vertical and horizontal transmission of the element in bacterial populations.

Prototypic Tn7 moves by a cut-and-paste mechanism where the element is completely excised from a donor site by a heteromeric transposase composed of TnsA and TnsB proteins (2, 3)(Figure 1A). The TnsB transposase is an RNaseH family protein that breaks the DNA at the 3’ ends of the element and directly joins these ends to the target DNA. TnsA is an endonuclease that is unique to Tn7 and Tn7-like elements that cleaves at the 5’ ends of the transposon to allow the element to be completely excised from the donor DNA (4–6). In one clade of Tn7-like elements, the TnsA and TnsB proteins are fused as a single transposase (7, 8). In some other clades of elements, TnsA and TnsB can be sporadically found fused. Regardless of the transposase configuration, transposase activity is only activated when dedicated target site selecting proteins or systems identify an appropriate target DNA and interact with the transposase through TnsC, a signaling protein belonging to the AAA+ family (9). Prototypic Tn7 and most Tn7-like elements recognize a preferred target in bacterial chromosomes called an attachment (*att*) site using TnsD, a TniQ-domain containing protein (10). Tn7 recognizes the *attTn7* site associated with the essential and highly conserved *glmS* gene, but TnsD family proteins evolved to recognize a variety of genes as *att* sites (also known as homing sites) across Tn7-like elements (Figure 1)(11–13).

**Figure 1:**
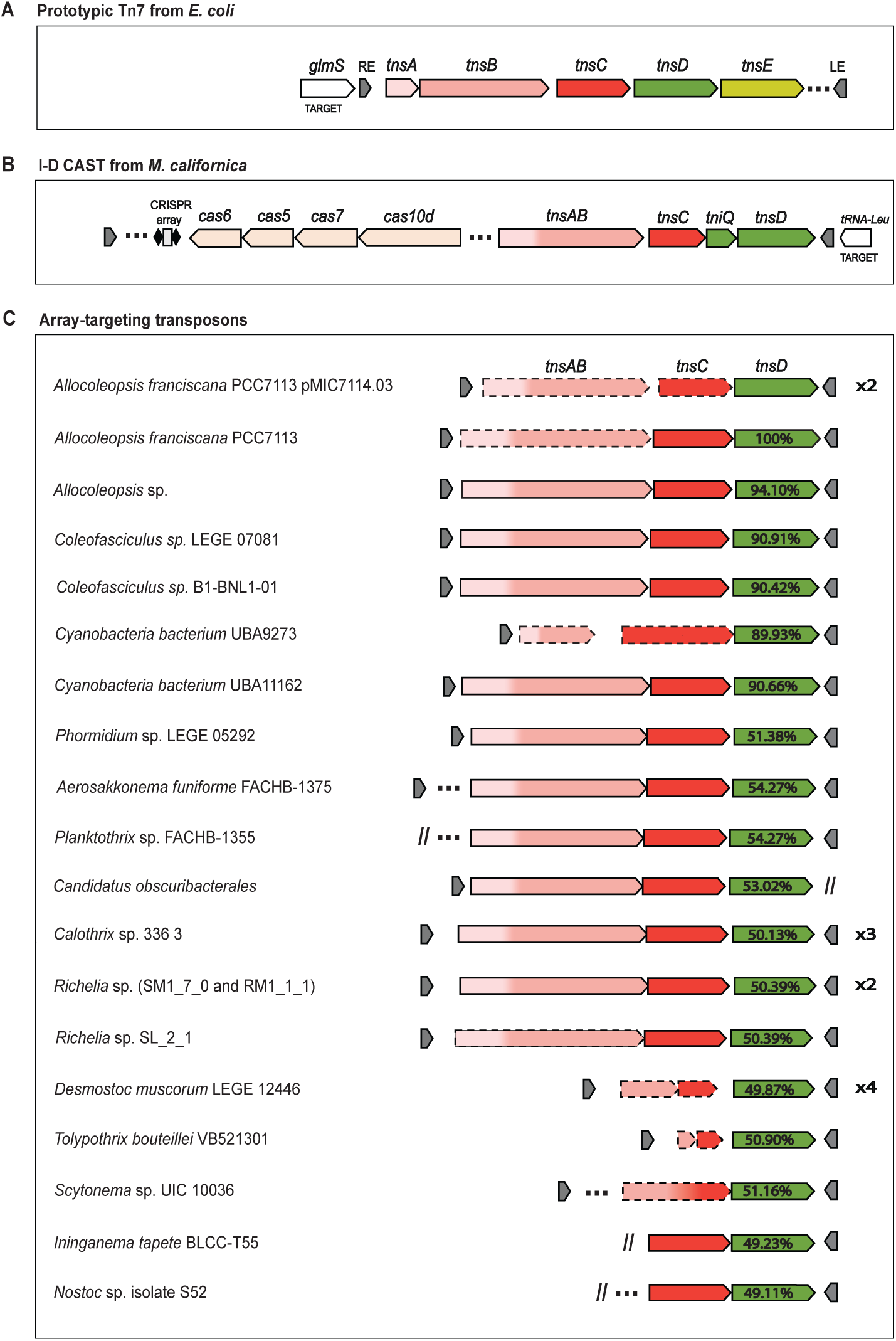
Family of Tn7-like transposons targeting CRISPR arrays. Graphic representation of prototypic Tn7 from *E. coli* (A), I-D CAST from *M. californica* (B), and all Tn7-like elements belonging to the array-targeting family (C). Each element within the array-targeting family is unique. The percent of identity of TnsD proteins from this group is shown within *tnsD* block arrow relative to TnsD from the element found in plasmid PMIC7114.03 of *A. franciscana* PCC7113, used as a reference. The cyanobacterial species in which each element was discovered is indicated on the right side of the graph. If an element was found multiple times, this is noted on the left side of the transposon. “RE” and “LE” denote Right End and Left End, respectively. Dash-lined arrows indicate genes containing internal stop codons or deletions, while three points represent genes not involved in transposition. Double diagonal lines signify the end of a contig.

In an alternate targeting pathway, Tn7 preferentially directs transposition into mobile plasmids and bacteriophages, using a different target site selecting protein TnsE that recognizes features of DNA replication (14–17). Diverse modalities have evolved to recognize mobile plasmids in other Tn7-like elements including multiple independent cooption events of diverse CRISPR-Cas system for natural guide RNA-directed transposition (18). These CRISPR Associated Transposons (CASTs) coopted various types of CRISPR systems, such as Type I-F, Type I-B, Type I-D and a Type V (V-K) system (7, 8, 19–23). As part of the adaptation process the CRISPR systems lost the capacity to degrade DNA (interference). In CAST systems the Cas components interface with the transposition machinery via a TniQ protein. The coopted CRISPR-Cas components are encoded between the cis-acting ends of the transposon and mobilize with the element as it moves to new locations.

Here we describe a subgroup of cyanobacterial Tn7-like transposons that adapted with CRISPR-Cas systems in a novel way, directing transposition into CRISPR arrays likely by recognizing CRISPR repeats as *att* sites. Unlike CAST elements, here the CRISPR-Cas machinery is not mobilized as part of the transposon machinery, but instead CRISPR arrays are directly targeted by the elements. Array-targeting and tRNA-targeting transposons (the ones that utilize TnsD protein) likely share a recent common ancestor as both types of elements are phylogenetically related and recognize a DNA sequence shared between the targets that are recognized. CRISPR-Cas systems have been shown to be common in bacterial chromosomes and on mobile plasmids (24). Therefore, the ability to target CRISPR arrays would provide the benefits of a safe chromosomal site favoring vertical transmission, and a feature common on mobile plasmids to favor horizontal transmission. The discovery of Tn7-like transposons targeting CRISPR arrays represents an additional novel coadaptation between mobile genetic elements and bacterial defense systems.

## Results

### A Tn7-like transposon targeting CRISPR arrays

Multiple laboratories have searched DNA databases for associations between Tn7-like transposons and CRISPR-Cas systems. These efforts have revealed numerous functional associations where CRISPR components have been naturally coopted for guide RNA (gRNA)-directed transposition. In CASTs, the system is encoded between the right and left ends of the transposon (Figure 1B). Interestingly, as part of our work we identified a family of Tn7-like elements that were also linked to CRISPR-Cas machinery, though the relevant genes were not encoded within the transposon itself. Instead, they were consistently found proximal to the element, outside of the transposon’s boundaries. Further analysis suggested that these elements specifically targeted the associated CRISPR arrays (see below).

In one example in the plasmid pMIC7113.03 from the filamentous cyanobacterium *Allocoleopsis franciscana* PCC7113 we found two identical copies of a Tn7-like transposon situated within distinct CRISPR arrays (Figure 1C). The element encodes the characteristic proteins found in transposons of this family, TnsA and TnsB (fused), TnsC, and TnsD. A near-identical copy of the element was found in the chromosome of *A. franciscana*, but not adjacent to a CRISPR array. The chromosomally encoded version is likely non-functional due to nonsense mutations in the transposase gene. The plasmid version shows deletions in *tnsAB* and *tnsC* genes thus it is probably also inactive.

However, we could identify a highly similar element (93.4 % nucleotide identity when compared to the chromosomal version) in another *Allocolepsis* sp. genome with predicted fully functional genes (Accession NZ_DATNPA010000798) suggesting this element is active in cyanobacterial populations (Figure 1C).

### A family of array-targeting transposons

Homology searches with the BLAST algorithm using the amino acid sequence of TnsD from *A. franciscana* identified 16 unique closely related homologs with ∼50-90% identity (Figure 1C). Subsequent analysis revealed their association with 19 unique transposons possessing similar characteristics to the elements discovered in *A. franciscana* and *Allocolepsis sp.* (Figure 1C). Shared characteristics included fused TnsA and TnsB proteins, something common to the majority of elements in this clade, a TnsC AAA+ protein, and a single TnsD-like target selection protein. The TnsD homologs all maintained the CCCH zinc finger motif predicted to be involved in DNA binding (25) (Supplemental Figure 1).

Notably, examination of the transposon context revealed that ∼82% (18 out of 22) of the elements whose boundaries could be examined are also situated within CRISPR arrays, suggesting CRISPR arrays are preferentially targeted by this group. Many of the elements were identified on small DNA contigs, so information outside the CRISPR array could not be assessed in all cases. The transposons whose context could be examined included insertions into orphan arrays and into arrays directly proximal to Cas encoded proteins (i.e. full contiguous systems). Besides the element found in *A. franciscana*, some other members of the family also appear to be recently active as we found multiple examples of identical or near identical copies in the same genome, in different strains of the same species, or in closely related species (Figure 1C). For instance, elements from *Calothrix* sp 336 3 and *Desmonostoc muscorum* LEGE 12446 are found three and four times in their respective genomes and near identical elements are found in three different *Richelia* strains (Supplementary Table 1). Interestingly, elements in *Calothrix* sp. 336/3 were identified twice within CRISPR arrays and a third time in a non-CRISPR array target, specifically at the end of a gene coding for a tyrosine phosphatase family protein, indicating that transposition is not limited to CRISPR arrays (see below).

The array-targeting elements appear to be minimal transposons that do not carry cargo genes. Cargo genes encode proteins that are not directly involved in the transposition process, but instead provide beneficial functions to the host, such as phage defense systems or antibiotic resistance determinates (26, 27). Only three members of the family (identified in *Planktothrix*, *Scytonema*, *Nostoc sp*.) appeared to have genetic information beyond the *tnsAB, tnsC,* and *tnsD* genes and these appeared to be different kinds of independent competing mobile genetic elements that integrated within the transposons disrupting core transposition genes.

Bacterial genomes often encode Tn7-like elements that are predicted to be inactive because genes encoding transposition proteins are partially degraded. However, we found examples where elements that were predicted to be inactive were found in multiple identical copies, suggesting they were in-fact mobile. For example, as in the plasmid of *A. franciscana*, where there are identical copies of an element with *tnsAB* and *tnsC* genes seemingly degraded. That is also the case with the element found in *Desmonostoc muscorum* LEGE 12446 where a deletion fused the *tnsAB* and *tnsC* open reading frames into a likely nonfunctional single 270-amino acid protein, but four identical copies of the same transposon were found in the genome. Similarly, the elements from *Tolypothrix bouteillei* VB521301 and *Scytonema* sp. UIC 10036 exhibit long deletions in their *tnsAB* and *tnsC* proteins while *tnsD* and the transposon ends remain intact. It is possible that these partially degraded elements could be mobilized in trans if the cis-acting transposon ends recognized by the transposase remain intact and are recognized by protein components expressed from another element in the genome. It is also possible that all these elements required trans-complementation with a complete or complementary version of the element that is no longer found in the genome.

### Array-targeting elements are closely related to tRNA-targeting elements in Cyanobacteria

To analyze the phylogenetic context of the array-targeting elements, we carried out a HMMER search of all available protein sequences from Cyanobacteria in NCBI using the Pfam TniQ profile (PF06527). A similarity tree of nonredundant TnsD/TniQ proteins revealed that the array-targeting transposons are phylogenetically close to TnsD-mediated tRNA-targeting transposons in this phylum (Figure 2, 3 and 4). Elements that recognize tRNA genes identified in this search include examples that had been previously reported and experimentally validated like I-D and I-B2 CAST (7, 22). tRNA genes are known hotspots for mobile element integration in bacteria including bacteriophages and integrating conjugating elements possibly because such genes are highly conserved (28–30). Of note, TnsD from *A. franciscana* (AfTnsD) shares 42.3% of homology with TnsD from I-D CAST found in *Myxacorys californica* (McTnsD) where tRNA-Leu targeting has been confirmed in the heterologous *E. coli* host (Figure 1, 3) (12, 31). We noted that a distinct structural difference that co-occurs with the change in *att* site used between these two elements is a C-terminal motif found in the tRNA-targeting TnsD proteins that is absent in the array-targeting group (Figure 3 and 5). AlphaFold3-generated structures suggest that the C-terminal region absent in AfTnsD representative corresponds to a helix-turn-helix domain (HTH) (Figure 5). Interestingly, the C-terminal region of TnsD is known to be involved in target-specific DNA binding in canonical Tn7 (25) and predicted to have a similar function in other TnsD-like proteins (32).

**Figure 2:**
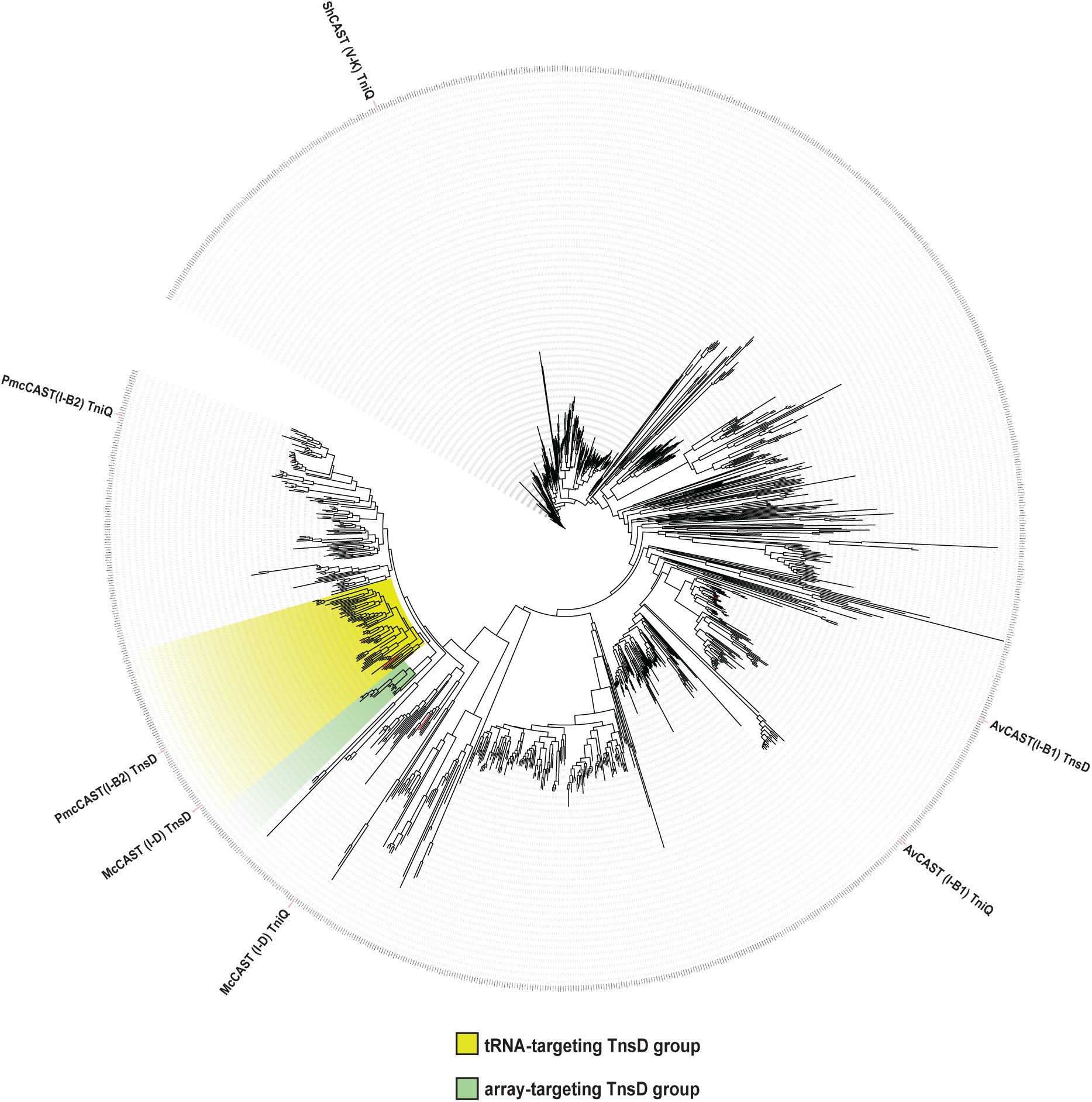
TniQ similarity tree from Cyanobacteria. A TniQ similarity tree constructed using all TniQ proteins obtained from cyanobacterial genomes available at NCBI via *hmmsearch* (TniQ hmm profile PF06527.hmm). The clade containing TnsD proteins predominantly recognizing tRNA genes as targets is highlighted in yellow, while the clade of array-targeting TnsD proteins is highlighted in green. Proteins associated with known CAST systems are marked.

**Figure 3:**
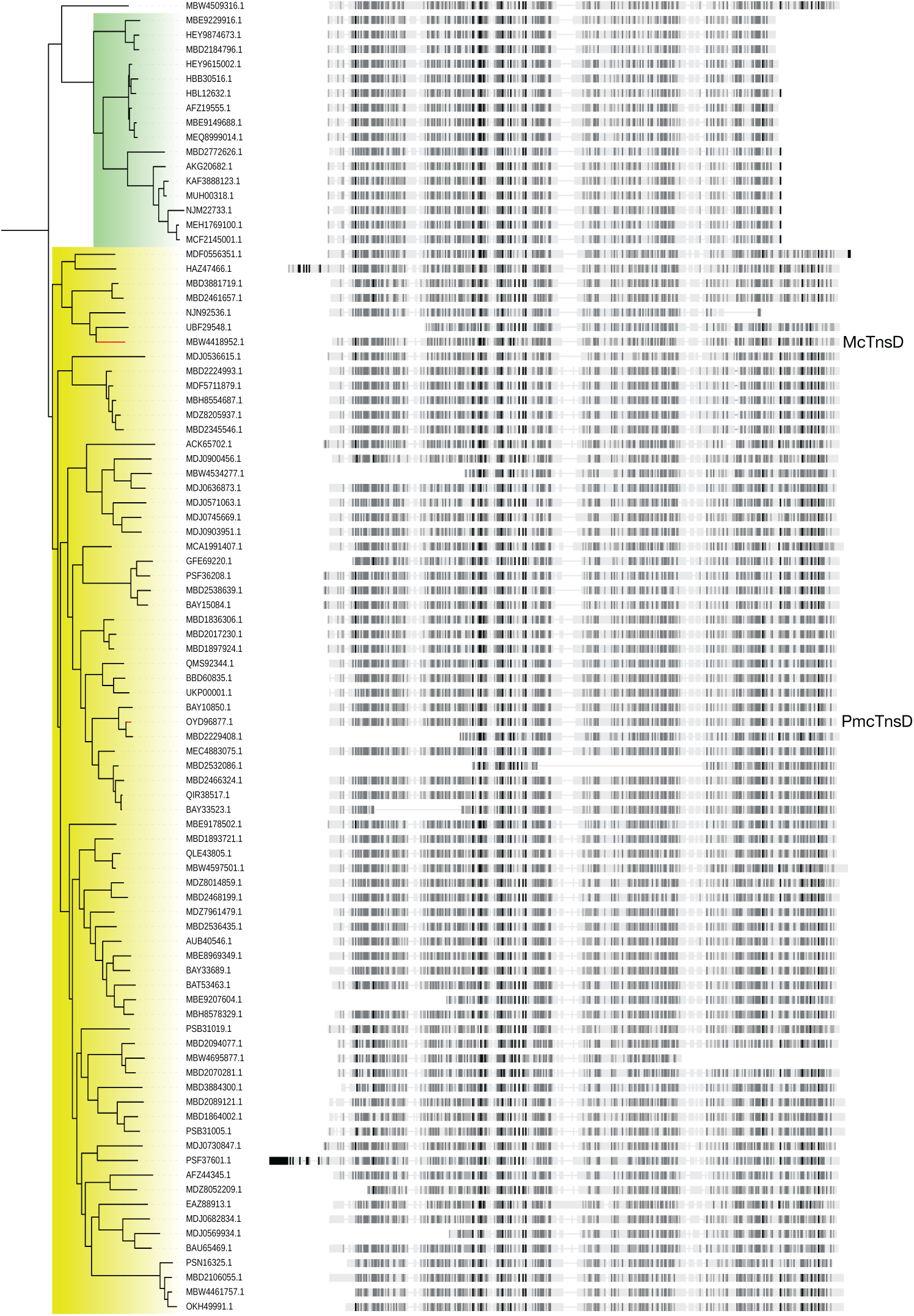
TnsD Proteins from the “yellow” and “green” clades of TniQ Tree. Left: Rooted view of the yellow- and green-highlighted tree clades from Figure 2, including accession numbers. Right: Graphic representation of the corresponding protein alignment. The darker the color, the higher the sequence homology.

### Sequences common to tRNA genes and CRISPR arrays provide a clue for how array-targeting elements might have evolved to recognize a different target

DNA sequences recognized by TnsD are known to occur at a fixed distance from the point of integration with Tn7 and Tn7-like transposons. Aligning the DNA sequences that flank the ends of the tRNA- and array-targeting elements revealed a conserved DNA motif 40 to 50 bp away from the transposons left ends (Figure 6A-B). Of further interest, this sequence mapped to a position that is conserved across both the tRNA genes and the repeats from the CRISPR arrays where array-targeting elements are found. Although not all array-targeting elements are found adjacent to a CRISPR array (Figure 4, examples indicated in Figure 6A by blue dots), their predicted targets still maintained this sequence match, suggesting it also accounted for integration events outside of CRISPR arrays (See discussion).

**Figure 4:**
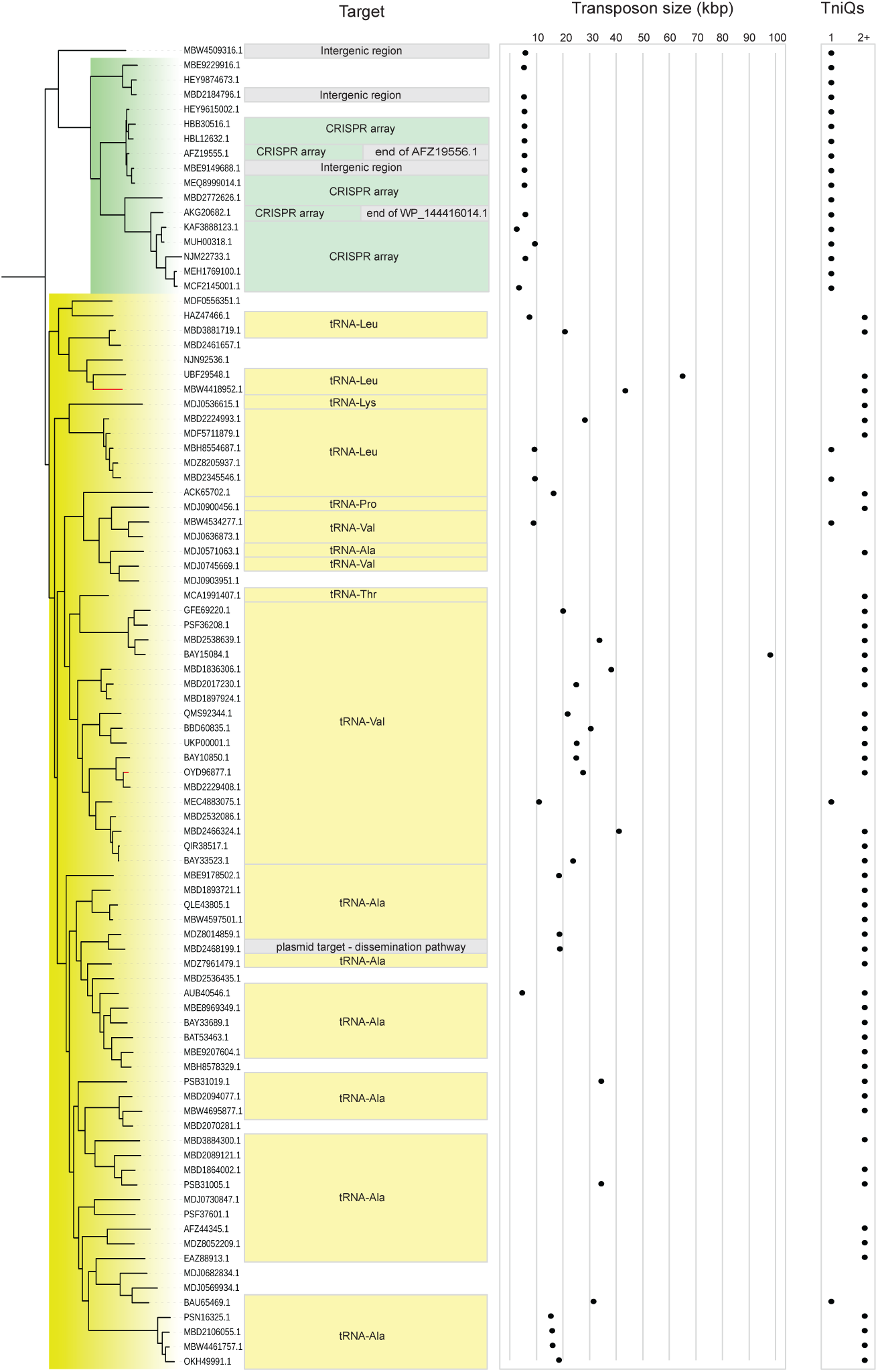
Characteristics of the transposons associated with CRISPR arrays and tRNA genes. A similarity tree is shown for TniQ (rooted). The accession number, location (target), approximate size, and the number of TniQs encoded in the element are indicated. Left: Rooted view of the yellow-highlighted tree branch from Figure 1. The size is provided only for transposons where both ends could be detected.

**Figure 5.**
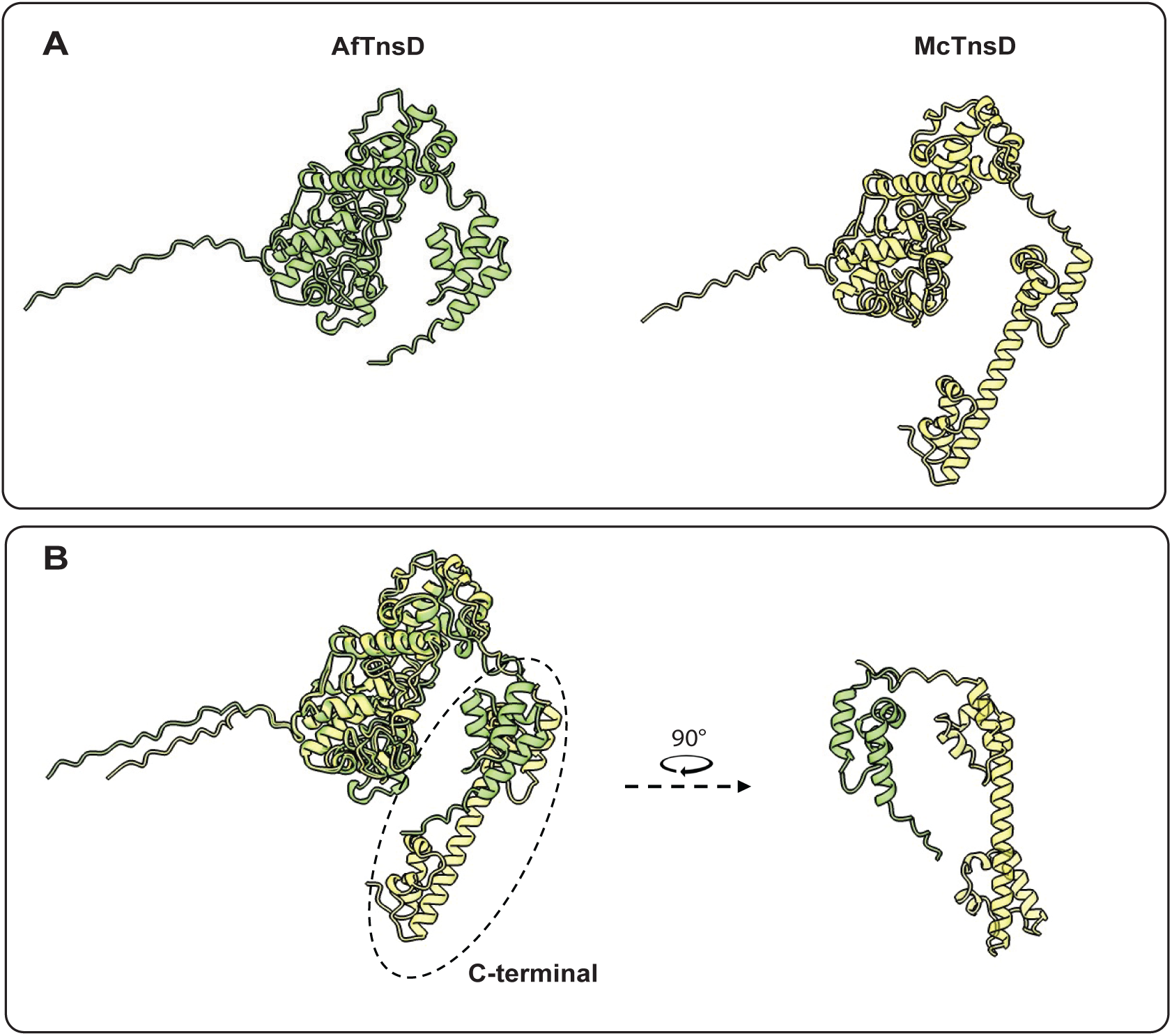
Aplphafold3 predicted three-dimensional structures of representative TnsD proteins from the tRNA- and array-targeting groups of transposons. (A) Yellow: McTnsD (I-D CAST from *M. californica)*. Green: AfTnsD (array-targeting from *A. franciscana*). (B) Superimposed structures of both proteins using the matchmaker function from ChimeraX.

**Figure 6.**
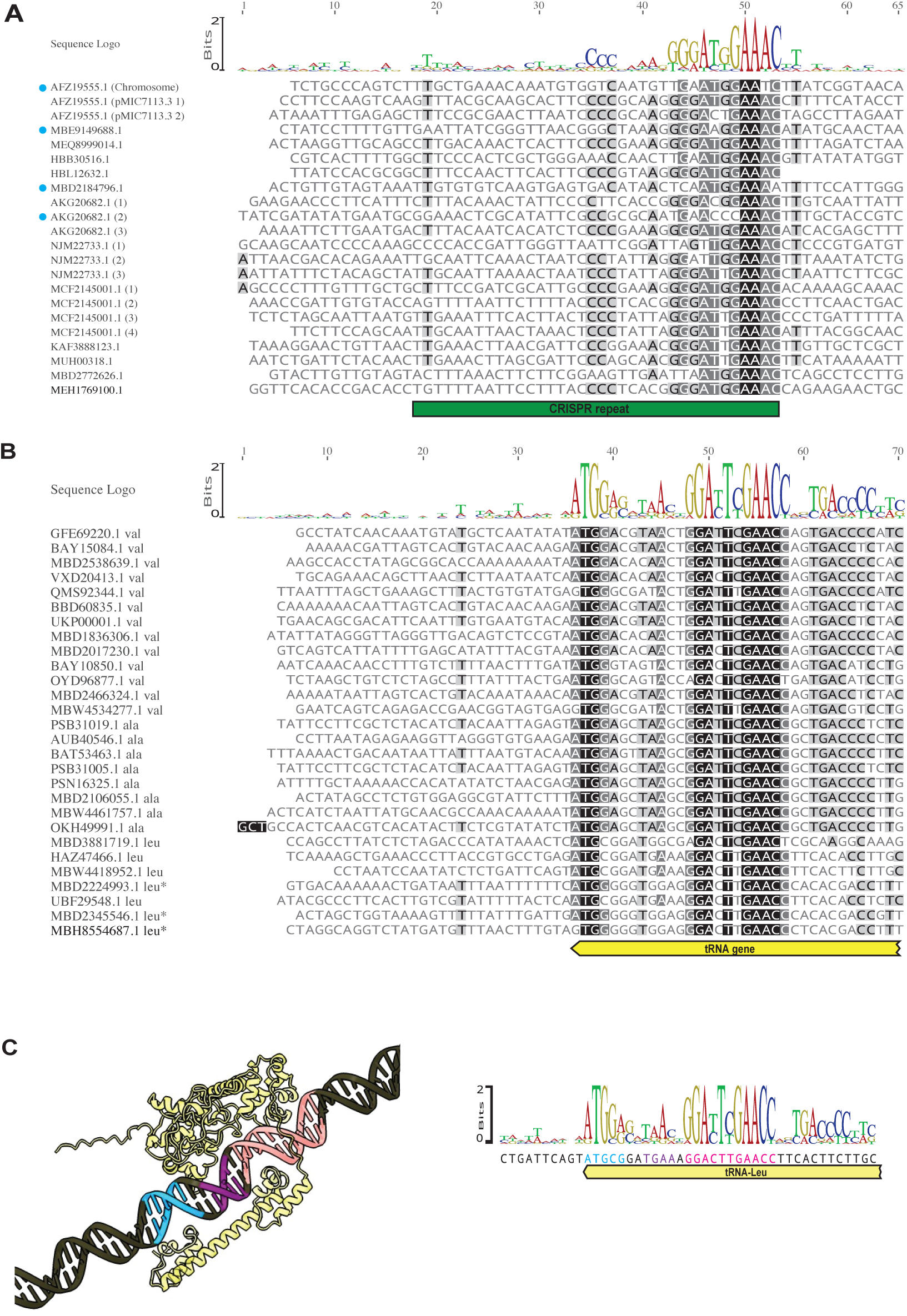
Predicted attachment site of array-targeting transposons. The figure shows the alignment of the sequences immediately downstream the left end of transposons targeting (A) CRISPR arrays and (B) tRNA genes. The sequence logo is shown on the top of each alignment. The darker the background of a nucleotide, the higher the sequence homology. The region corresponding to the CRISPR repeat is marked with a green box at the bottom of the alignment. Elements whose target is not a CRISPR repeat are marked in the alignment with a blue dot. The region corresponding to the end of the tRNA gene is marked with a yellow box at the bottom of the alignment. The shortened name of the specific tRNA gene targeted (Val, Ala, Leu or Leu*) is written next to the accession number. Only downstream sequences of elements where both ends were detected were used for the alignment. (C) Alphafold3-generated structure of McTnsD-target DNA.

AlphaFold3 accurately modeled the McTnsD-target DNA complex (Figure 6C), but failed to model the binding of AfTnsD to its predicted target. According to the model, four different regions of McTnsD interact with the end of tRNA-Leu gene, coinciding the recognized sequences with conserved regions across tRNA genes (colored in blue, purple and pink in Figure 6C). The conserved region colored in pink is, as mentioned above, also conserved in CRISPR arrays. However, the conserved region colored in blue, recognized by the HTH domain at the C-terminal of McTnsD (that is absent in AfTnsD) does not have a parallel in CRISPR arrays. The predicted structure is in agreement with the recently published CryoEM structure of PmcTnsD (from I-B2 CAST) bound to its tRNA-Val target (33).

Using CRISPR-Finder, we determined the potential orientation of each array relative to the orientation of the transposon. Although the leader region was not identified by this software, the potential orientation of the array was inferred from the A-T content of the array-flanking regions. The analysis revealed that all the insertions occurred in the same orientation with CRISPR arrays. Although the orientation of integration events was conserved, we did not observe any specific preference for transposons to bias transposition to any particular repeat position (i.e., near the leader, in the middle, or near the end). We suspect that integration into the CRISPR array does not compromise its ability to function. This is based on the observation that even when integration events occurred early in the array, there was no obvious degradation of the downstream repeats.

### Genetic evidence of specialization within individual subgroups of array-targeting elements

Among the array-targeting family, three subgroups can be identified based on TnsD similarity (indicated in orange, blue and green in Figure 7). When analyzed closely, other distinctive features were found in each of the subgroups. The differences include slight changes that were conserved in the subgroups like distinct lengths of TnsAB and TnsD proteins and variations in the sequences and lengths of the right and left ends.

**Figure 7.**
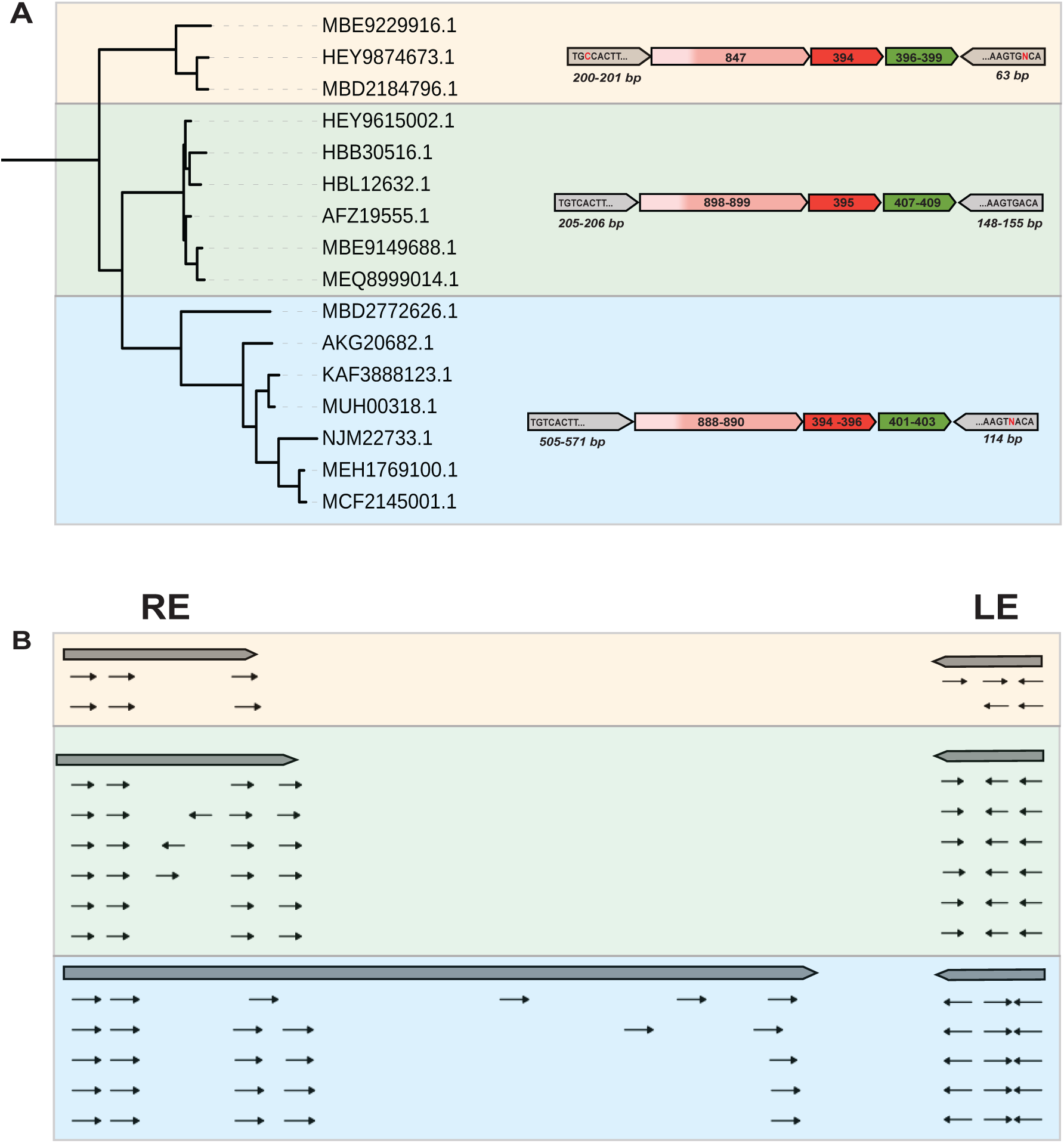
Signs of specialization within the array-targeting family. (A) TniQ similarity tree of the array-targeting clade. Three sub-groups were identified based on TniQ similarity. Additional differences were identified between the tree sub-groups and are graphically represented. Numeric values inside arrows represent amino-acid sizes while all other numeric values represent distance in base pairs. (B) Distribution of TnsB-binding sites across the right and left ends of transposons from elements indicated above.

Interestingly, the branch indicated in blue evolved an unusually large right end, averaging 500 bp (Figure 7A). These regions in Tn7-like transposons usually contain transposase binding sites for coordinating transposition or autoregulating the transposition genes (34). Curiously, elements from the blue branch also exhibit very distinctive patterns of TnsB-binding site distribution and orientation in both right and left ends, when compared to elements from the other two branches and I-D CAST from *M. californica* (Figure 7B) (7). Of note, the majority of elements that were found multiple times in the same or closely related genomes (an indirect piece of evidence supporting higher transposition activity) lie in this subgroup. While these differences support some kind of specialization within the array-targeting family, the mechanistic underpinning of these differences remain unclear.

### The array-targeting elements could not be established in the heterologous *E. coli* host

We attempted to establish the array-targeting systems from *A. franciscana* (Figure 7, green sub-group) and *Calothrix* sp. (Figure 7, blue sub-group) in an *E. coli* host using a mating-out assay (see details in Methods). As a positive control, a transposon with a similar TnsD protein that had previously been established in the heterologous *E. coli* host was included, the TnsABCD system from *M. californica*. None of the array-targeting elements that were evaluated showed targeted or random transposition activity under the tested conditions (Supplementary Figure 2, data not shown for *Calothrix* sp.). We also tested heterologous complementation between array-targeting and tRNA-targeting elements by shuffling their core proteins and targets in different combinations but none of them showed activity. In addition, we evaluated whether the very C-terminal of TnsD (the one that array-targeting elements lack) is crucial for target recognition by deleting this fragment in McTnsD. Deletion of this domain in Mc-TnsD caused a total loss of transposition activity.

## Discussion

We discovered a family of cyanobacterial Tn7-like transposons that are predicted to use the repeats in CRISPR arrays as *att* sites. Phylogenetically, these elements appear to share a common ancestor with tRNA-targeting Tn7-like transposons. Given that CRISPR-Cas systems can be common on both chromosomes and mobile plasmids, the ability to target CRISPR repeats would provide a stable reservoir for these elements in chromosomes and a means to facilitate transfer to new bacterial hosts using a single target capture system. While these are minimal elements that do not carry genetic cargo, a feature generally found with Tn7-like elements, they could play a role with mobilizing CRISPR-Cas systems between bacteria if excised as composite transposons.

### Array-targeting transposons are minimal Tn7-like elements

Unlike most Tn7-like elements, array-targeting transposons do not accumulate cargo genes that would provide accessory functions to the host or to benefit the transposon directly. This contrasts with what is found with the closely related tRNA targeting elements, which usually carry two TniQ domain-containing proteins and consistently accumulate cargo genes, resulting in a much larger transposon size (Figure 4). In this way, the array-targeting elements appear to behave more like classic insertion sequences, the smallest transposable elements found in prokaryotic genomes, that usually encode only a transposase gene and are found multiple times in the same genome (35). We suspect that array-targeting transposons do not negatively impact the functioning of the CRISPR-Cas systems where they reside. Residing silently in the CRISPR array could be important and preclude the elements from accumulating additional genes. Given that CRISPR arrays often are found to flank both sides of sets of Cas genes, having array-targeting elements in arrays that bracket an entire CRISPR-Cas system could allow the system to be mobilized to new hosts. Interestingly, this configuration was found in the filamentous cyanobacterium *Allocoleopsis franciscana* PCC7113 pMIC7113.03 plasmid. An analogous role was also suggested for site specific recombination systems that can mobilize CRISPR-Cas systems in cyanobacteria (36).

### Targeting CRISPR arrays would provide access to chromosome and plasmid targets

Prototypic Tn7 has complementary transposition targeting pathways. The TnsD protein allows the element to target the conserved *attTn7* site downstream of the *glmS* gene. The TnsE pathway recognizes features of conjugal DNA replication to facilitate transfer between bacterial hosts. The broader family of Tn7-like elements has continually re-evolved this two-pathway lifestyle, including all CAST elements (18). We suggest that because CRISPR-Cas systems are commonly found in the chromosome and encoded on plasmids in cyanobacteria, targeting arrays could allow a special benefit in this group. It would allow both a safe site in bacterial chromosomes and a means to facilitate transfer between bacterial hosts via plasmids. It has been reported that complete CRISPR-Cas systems and orphan arrays are widespread among filamentous cyanobacteria (36).

### Integration into CRISPR arrays and defense system function

We have no evidence that integration of minimal elements compromises the preCRISPR RNA from being transcribed or processed as functional RNA guides. Usually Tn7-like transposons avoid negative consequences to the host with their integration by recognizing essential genes but integrating outside the coding region of the gene. However, there are also examples where Tn7-like elements such as Tn6022 evolved specifically to inactivate a gene, the *comM* gene involved in natural competence (31). Interestingly, CAST elements independently evolved *comM* targeting and targeting directing transposition into other competence genes indicating this a general property of Tn7-like elements, something also found with other integrating elements (12, 31).

There is precedent in another system where CRISPR repeats are targeted with a bacteriophage, where the downstream guides are inactivated. Bacteriophage ΦAP1.1 uses host CRISPR repeats as an integration site, and in this case integration neutralizes part of the immune response; all spacers downstream the lysogen are not transcribed and the spacer immediately upstream of the interrupted repeat is not processed into a functional crRNA guide (37). In contrast to the ΦAP1.1 phage, array-targeting transposons do not disrupt the repeat structure as they insert in the spacer area. This together with the unusually small size of the transposons from this family could indicate that the transcription and processing of the CRISPR arrays is not affected, which is more consistent with the way Tn7-like elements function, remaining neutral to host functions.

### Array-targeting transposons may rely on a host factor or factors

We were unable to reconstitute array-targeting elements in *E. coli* using an *in vivo* assay that has been successfully employed in our lab to test a wide variety of Tn7-like transposons (7, 11). These elements do not seem different *a priori* from their tRNA-targeting relatives that have been experimentally validated in other work (7, 22), except for the absence of a ∼60 amino acid C-terminal region in TnsD proteins from array-targeting transposons. As showed in the Results section, this region is predicted to be part of the DNA binding domain in tRNA-targeting transposons, and we proved it is essential for TnsD-mediated transposition in McCAST. It is possible that a DNA-binding deficient TnsD relies on a host factor or factors to assist with target DNA recognition. For canonical Tn7, two proteins, LCP and L29, are known host factors responsible for high frequency transposition into the *attTn7* site at *glmS* (38). Cyanobacteria and gamma Proteobacteria are highly diverged branches of bacteria, thus potential host factors might be absent or too different in the heterologous host to function.

### Concluding remarks

The work presented here provides a glimpse into a novel way that Tn7-like elements have adapted in Cyanobacteria. Bioinformatics work in cyanobacteria is challenging because of the limited number of genomes that are available in this exciting group of bacteria. Moreover, there is a limited number of fully sequenced and closed genomes available to support a more thorough analysis of array-targeting transposons. The majority of the elements reported here are found in fragmented contigs complicating efforts to fully understand the genomic context of these elements on a broader scale. Within this complexity there is likely a wealth of untapped genetic variability awaiting discovery with more cyanobacterial genomes.

## Methods

### Computational methods

Annotated genomic sequences and feature tables of Cyanobacteria were downloaded from National Center for Biotechnology Information (NCBI) FTP site. There were 4094 annotated genomes when sequences were downloaded for our analysis (August 2024). The HMM profile associated with TniQ (PF06527) downloaded from the European Bioinformatics Institute (EMBL-EBI) Pfam database, was used for detecting homologs with *hmmsearch* (HMMER3). Proteins with more than 90% identity were clustered together using cd-hit. Entries marked as ‘partial’ were removed, reducing the final protein list to 1095 sequences (Supplementary Table 2). The curated protein list was aligned using MUSCLE (39) and a similarity tree was constructed using FastTree (40). The tree was visualized in iTOL (41).

Alignments of the right and left ends of array-targeting and tRNA-targeting transposons shown in Figure 6 and of TnsD proteins shown in Figure 3 were made using Clustal Omega accessed from Geneious Prime 2024.0.7. For tRNA-targeting examples in Figure 6B, the analysis only included elements whose right and left ends were identified. Three-dimensional structures of TnsD proteins and TnsD-DNA complex were obtained from AlphaFold3 (42) and visualized in ChimeraX (43).

### Bacterial strains and plasmids

Found in Supplementary Table 3.

### Mating-out assay

The assay monitors the movement of the mini-element (the transposon right and eft ends flanking a Kanamycin - Kan-resistance gene) from the chromosome of the donor BW27783 (44) cells to a fertility (F) plasmid (carrying a Spectinomycin - Spec-resistance gene) containing the putative DNA target sequence recognized by the transposon. Transposition proteins are expressed from inducible plasmid vectors in the donor cells. After protein induction, the F plasmid is mated into recipient CW51 (45) cells (Nalidixic acid -Nal- and Rifampicin - Rif-resistant) and the cell mix is plated in the appropriate antibiotics. The transposition frequency is determined as the percentage of recipient cells with the transposon marker from the total of recipient cells that received the F plasmid.

In more detail, overnight cultures of donor and recipient cells were made from single colonies at 37°C in 5 ml of Luria-Bertani (LB) broth supplemented with Chloramphenicol - Cam-(30 µg/mL), Carbenicillin - Carb-(50 µg/mL) and glucose (0.2%)(46). Overnight cultures were washed with plain LB twice and sub-cultured 1:50 in fresh minimal M9 media supplemented with maltose (0.2%) as carbon source, Cam, Carb, arabinose (0.2%) and IPTG (100 µM). During induction, donor strains were grown overnight with aeration at 30°C. In parallel, an overnight culture of the recipient cells was made from a single colony at 37°C in 5 ml of LB. On the next day, donor and recipient cells were washed with LB media, mixed in a ratio of 1:10 (donor: recipient) and grown at 37°C on a culture wheel turning at the lowest speed to facilitate mating. The cell mix was serial diluted, and the correct dilutions were plated in LB agar containing the appropriate antibiotics. The cells were simultaneously plated in Spec (100 µg/mL), Nal (20 µg/mL), Rif (100 µg/mL) LB agar plates to calculate the total number of transconjugants and in Spec, Nal, Rif, Kan (50 µg/mL) LB agar plates to calculate the number transconjugants that received KanR gene via transposition.

## Additional material

Supplementary Figures

Supplementary Table 1 - Summary of array-targeting transposons

Supplementary Table 2 - Summary of TniQ proteins from Cyanobacteria

Supplementary Table 3 - Bacterial strains, plasmids and primers

## Acknowledgements

Michael Petassi for rationalizing the first ideas and designing the initial vectors of the project. Shan Chi (Popo) Hsieh, Saadlee Shehereen and Jordan Thesier for providing help with the construction and optimization of the TniQ similarity tree.

## Author details

Department of Microbiology, Cornell University, Ithaca, NY 14853, USA

## Authors’ contributions

LCM and JEP designed the experiments. LCM performed the experiments. LCM and JEP wrote the manuscript.

## Competing interest

The authors declare they have no competing interests

---

**Supplementary Figure 1.**
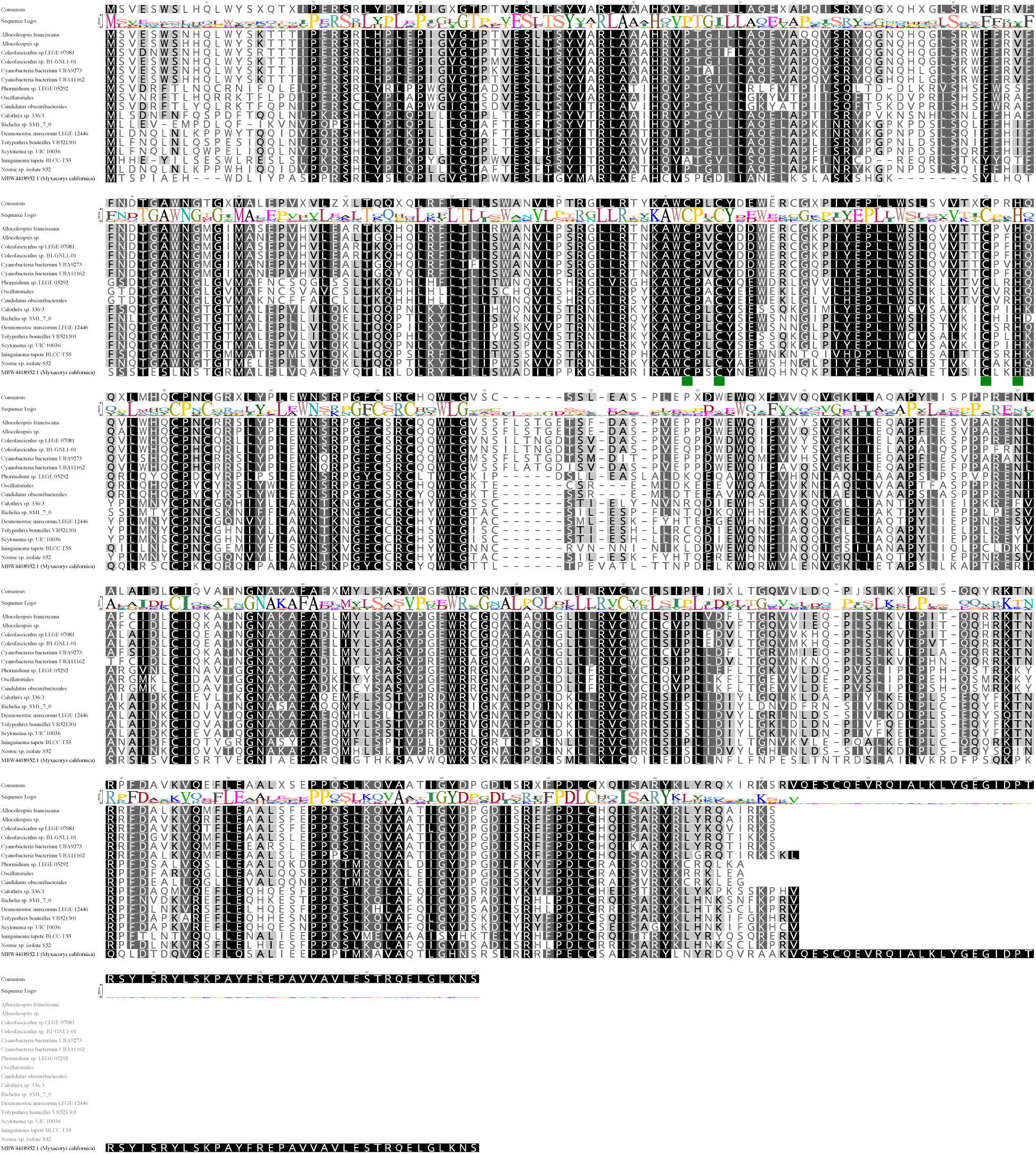
Alignment of TnsD proteins from the array-targeting group and TnsD from I-D CAST showing the conserved CCCH motif characteristic of TnsD-like proteins

**Supplementary Figure 2.**
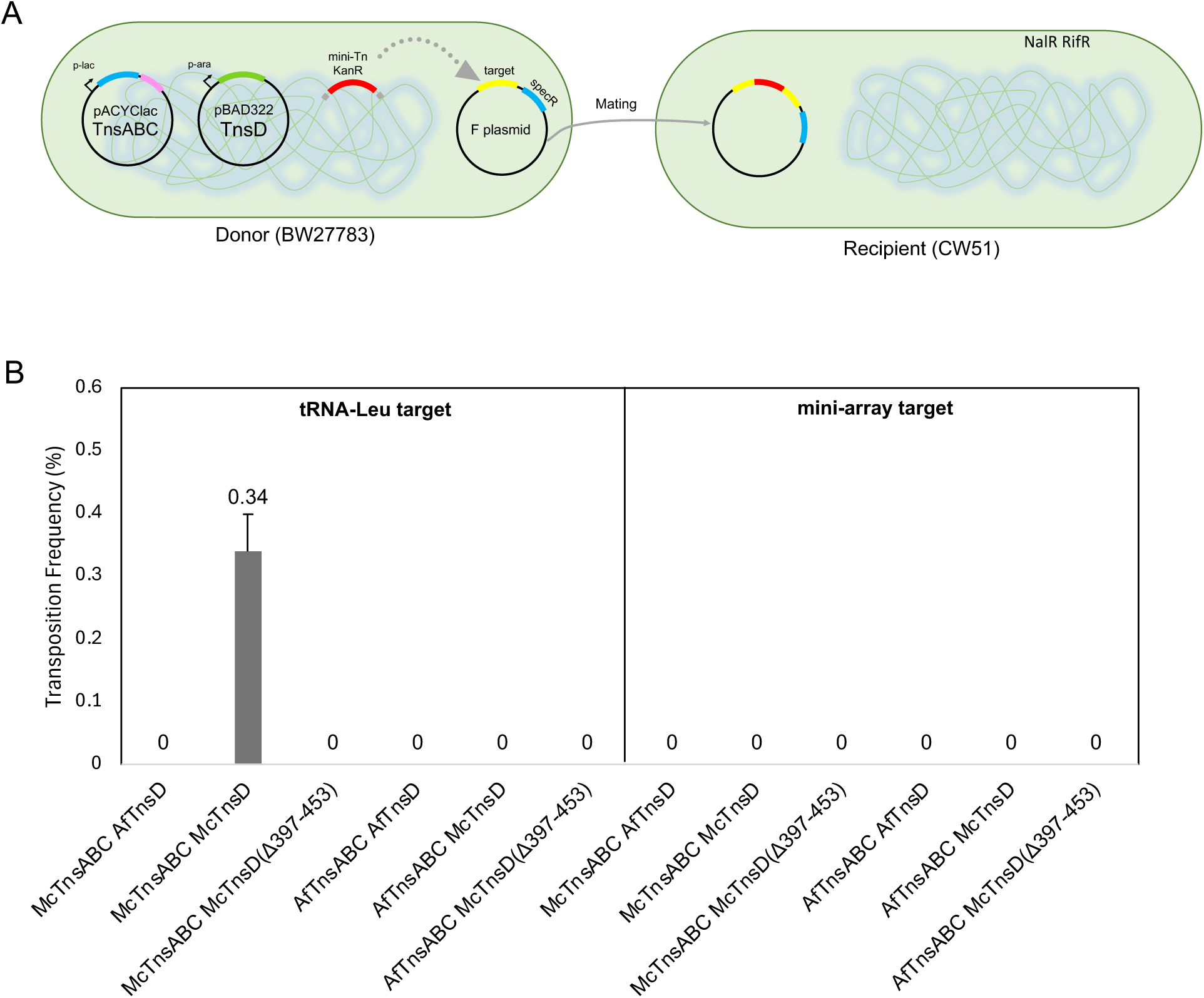
Testing functionality of the predicted array-targeting transposon from *A. franciscana*. (A) Diagram of the mating-out assay (see details in Methods). (B) Transposition frequency (%) for different combinations of transposition components from the previously tested tRNA targeting element from *M. californica* and the predicted array targeting element from *A. franciscana*.

## Notes

### Competing Interest Statement

The authors have declared no competing interest.

## References

1. J. E. Peters, Tn7. Microbiol Spectr 2, 1–20 (2014).

2. R. J. Bainton, K. M. Kubo, J.-N. Feng, N. L. Craig, Tn7 transposition: target DNA recognition is mediated by multiple Tn7-encoded proteins in a purified in vitro system. Cell 72, 931–943 (1993).

3. R. Bainton, P. Gamas, N. L. Craig, Tn7 transposition in vitro proceeds through an excised transposon intermediate generated by staggered breaks in DNA. Cell 65, 805–816 (1991).

4. R. Sarnovsky, E. W. May, N. L. Craig, The Tn7 transposase is a heteromeric complex in which DNA breakage and joining activities are distributed between different gene products. EMBO J. 15, 6348–6361 (1996).

5. E. W. May, N. L. Craig, Switching from cut-and-paste to replicate Tn7 transposition. Science 272, 401–404 (1996).

6. D. R. Ronning et al., The carboxy-terminal portion of TnsC activates the Tn7 transposase through a specific interaction with TnsA. Embo J 23, 2972–2981 (2004).

7. S. C. Hsieh, J. E. Peters, Discovery and characterization of novel type I-D CRISPR-guided transposons identified among diverse Tn7-like elements in cyanobacteria. Nucleic Acids Res 51, 765–782 (2023).

8. J. E. Peters, K. S. Makarova, S. Shmakov, E. V. Koonin, Recruitment of CRISPR-Cas systems by Tn7-like transposons. Proceedings of the Na@onal Academy of Sciences 114, E7358 (2017).

9. Z. Li, N. L. Craig, J. E. Peters, “Transposon Tn7” in Bacterial Integrative Mobile Genetic Elements, A. P. Roberts, P. Mullany, Eds. (Landes Bioscience, 2013), pp. 1–32.

10. R. L. McKown, K. A. Orle, T. Chen, N. L. Craig, Sequence requirements of Escherichia coli afTn7, a specific site of transposon Tn7 insertion. J Bacterial 170, 352–358 (1988).

11. M. T. Petassi, S. C. Hsieh, J. E. Peters, Guide RNA CategorizaIon Enables Target Site Choice in Tn7-CRISPR-Cas Transposons. Cell 183, 1757–1771 e1718 (2020).

12. G. Faure et al., Modularity and diversity of target selectors in Tn7 transposons. Mol Cell 83, 2122–2136.e2110 (2023).

13. A. Correa, 3rd et al., Novel mechanisms of diversity generation in Acinetobacter baumannii resistance islands driven by Tn7-like elements. Nucleic Acids Res 52, 3180–3198 (2024).

14. C. A. Wolkow, R. T. DeBoy, N. L. Craig, Conjugating plasmids are preferred targets for Tn7. Genes & Dev. 10, 2145–2157 (1996).

15. J. A. Finn, A. R. Parks, J. E. Peters, Transposon Tn7 directs transposition into the genome of filamentous bacteriophage M13 using the element-encoded TnsE protein. J Bacterial 189, 9122–9125 (2007).

16. A. R. Parks et al., Transposition into replicating DNA occurs through interaction with the processivity factor. Cell 138, 685–695 (2009).

17. Q. Shi et al., Conformational toggling controls target site choice for the heteromeric transposase element Tn7. Nucleic Acids Res 43, 10734–10745 (2015).

18. S. C. Hsieh, J. E. Peters, Natural and Engineered Guide RNA-directed transposition with CRISPR-Associated Tn7-like Transposons. Annu Rev Biochem 10.1146/annurev-biochem-030122-041908 (2024).

19. G. Faure et al., CRISPR-Cas in mobile genetic elements: counter-defence and beyond. Nat Rev Microbiol 17, 513–525 (2019).

20. S. E. Klompe, P. L. H. Vo, T. S. Halpin-Healy, S. H. Sternberg, Transposon-encoded CRISPR-Cas systems direct RNA-guided DNA integration. Nature 571, 219–225 (2019).

21. J. Strecker et al., RNA-guided DNA insertion with CRISPR-associated transposases. Science 365, 48–53 (2019).

22. M. Saito et al., Dual modes of CRISPR-associated transposon homing. Cell 184, 2441–2453 e2418 (2021).

23. S. E. Klompe, P. L. H. Vo, T. S. Halpin-Healy, S. H. Sternberg, Transposon-encoded CRISPR– Cas systems direct RNA-guided DNA integration. Nature 571, 219–225 (2019).

24. R. Pinilla-Redondo et al., CRISPR-Cas systems are widespread accessory elements across bacterial and archaeal plasmids. Nucleic Acids Res 50, 4315–4328 (2022).

25. R. Mitra, G. J. McKenzie, L. Yi, C. A. Lee, N. L. Craig, Characterization of the TnsD-afTn7 complex that promotes site-specific insertion of Tn7. Mob DNA 1, 18 (2010).

26. A. R. Parks, J. E. Peters, Tn7 elements: engendering diversity from chromosomes to episomes. Plasmid 61, 1–14 (2009).

27. S. Benler et al., Cargo Genes of Tn7-Like Transposons Comprise an Enormous Diversity of Defense Systems, Mobile Genetic Elements, and Antibiotic Resistance Genes. mBio 12, e0293821 (2021).

28. K. P. Williams, Integration sites for genetic elements in prokaryotic tRNA and tmRNA genes: sublocation preference of integrase subfamilies. Nucleic Acids Res 30, 866–875 (2002).

29. V. Burrus, M. K. Waldor, Shaping bacterial genomes with integrative and conjugative elements. Res Microbiol 155, 376–386 (2004).

30. B. Reinhold-Hurek, D. A. Shub, Self-splicing introns in tRNA genes of widely divergent bacteria. Nature 357, 173–176 (1992).

31. S. C. Hsieh, J. E. Peters, Discovery and characterization of novel type I-D CRISPR-guided transposons identified among diverse Tn7-like elements in cyanobacteria. Nucleic Acids Res 51, 765–782 (2022).

32. G. Faure et al., Modularity and diversity of target selectors in Tn7 transposons. Mol Cell 83, 2122–2136 e2110 (2023).

33. S. Wang, R. Siddique, M. C. Hall, P. A. Rice, L. Chang, Structure of TnsABCD transpososome reveals mechanisms of targeted DNA transposition. Cell 10.1016/j.cell.2024.09.023 (2024).

34. Z. Kaczmarska et al., Structural basis of transposon end recognition explains central features of Tn7 transposition systems. Mol Cell 10.1016/j.molcel.2022.05.005 (2022).

35. P. Siguier, E. Gourbeyre, M. Chandler, Bacterial insertion sequences: their genomic impact and diversity. FEMS Microbiol Rev 38, 865–891 (2014).

36. S. Hou et al., CRISPR-Cas systems in multicellular cyanobacteria. RNA Biol 16, 518–529 (2019).

37. A. Varble et al., Prophage integration into CRISPR loci enables evasion of antiviral immunity in Streptococcus pyogenes. Nat Microbiol 6, 1516–1525 (2021).

38. P. L. Sharpe, N. L. Craig, Host proteins can simulate Tn7 transposition: a novel role for the ribosomal protein L29 and the acyl carrier protein. EMBO J 17, 5822–5831 (1998).

39. R. C. Edgar, MUSCLE: multiple sequence alignment with high accuracy and high throughput. Nucleic Acids Res 32, 1792–1797 (2004).

40. M. N. Price, P. S. Dehal, A. P. Arkin, FastTree 2--approximately maximum-likelihood trees for large alignments. PLoS One 5, e9490 (2010).

41. I. Letunic, P. Bork, Interactive Tree of Life (iTOL) v6: recent updates to the phylogenetic tree display and annotation tool. Nucleic Acids Res 52, W78–W82 (2024).

42. J. Abramson et al., Accurate structure prediction of biomolecular interactions with AlphaFold 3. Nature 630, 493–500 (2024).

43. E. C. Meng et al., UCSF ChimeraX: Tools for structure building and analysis. Protein Sci 32, e4792 (2023).

44. A. Khlebnikov, K. A. Datsenko, T. Skaug, B. L. Wanner, J. D. Keasling, Homogeneous expression of the P(BAD) promoter in Escherichia coli by constitutive expression of the low-affinity high-capacity AraE transporter. Microbiology (Reading) 147, 3241–3247 (2001).

45. C. S. Waddell, N. L. Craig, Tn7 transposition: two transposition pathways directed by five Tn7-encoded genes. Genes Dev 2, 137–149 (1988).

46. J. E. Peters, “Gene transfer - Gram-negative bacteria” in Methods for General and Molecular Microbiology, C. A. Reddy et al., Eds. (ASM Press, Washington D. C., 2007), chap. 31, pp. 735–755.

